## Supplementary Figures for "Genome editing in animals with minimal PAM CRISPR-Cas9 enzymes" (from another version of this preprint: version 1, 2021-06-06)

#### Supplementary Figure legends

##### Supplementary Figure 1. Features of CRISPR-Cas9 systems

**a.** Scheme showing a gRNA (spacer in red, tail in blue) binding to the genomic target site (black) with the PAM sequence 5'-NGG (black, bold, and underlined), 5'-NGN (blue) and 5'-NAN (green) downstream for Cas9 WT-SpG-SpRY, SpG-SpRY or SpRY only, respectively. *In vitro* transcribed gRNAs were used for this study where a 5' GG requirement in the gRNA (between parenthesis) was needed for efficient transcription (Vejnar et al., 2016). Adapted from Moreno-Mateos et al., 2017.

**b.** Bar graphs showing number of targets of the indicated CRISPR endonucleases in the zebrafish and *C. elegans* genomes, separated in 5' UTRs, 3' UTRs, exons, genes, introns, and intergenic regions. AsCas12a: *Acidaminococcus* spp. Cas12a; ErCas12a: *Eubacterium rectale* Cas12a; LbCas12a: *Lachnospiraceae bacterium* Cas12a; SaCas9: *Staphylococcus aureus* Cas9; SpCas9: *Streptococcus pyogenes* Cas9; SpCas9 EQR: *S. pyogenes* Cas9 EQR mutant; SpCas9VQR: *S. pyogenes* Cas9 VQR mutant; SpCas9VRER: *S. pyogenes* Cas9 VRER mutant; SpGCas9: *S. pyogenes* Cas9 SpG variant; and SpRYCas9: *S. pyogenes* Cas9 SpRY variant.

##### Supplementary Figure 2. SpG and SpRY are efficient nucleases in zebrafish and *C. elegans*

**a.** Bar graphs showing the percentage of viability in the injected and non-injected embryos at 24 hpf. The embryos were injected with a mix containing 150 (1x) or 300 (2x) pg of mRNA coding for Cas9 WT (mCas9, grey), SpG (mSpG, blue), or SpRY (mSpRY, green) and 20 (1x), 120 (6x), or 240 (12x) pg of gRNA a or b targeting albino (**Fig. 1a**). Results are shown as averages ± standard error of the mean from two independent experiments with each gRNA (a or b) without significant differences between conditions (One-way ANOVA followed by Tukey's test for multiple comparisons).

**b-c.** Phenotypic evaluation of injected embryos with different amounts of mRNAs coding for Cas9 WT (mCas9), SpG (mSpG), or SpRY (mSpRY) and gRNA a (**b**) or b (**c**) targeting *slc45a2* (albino) showing different levels of mosaicism compared to the WT. Data from **Fig. 1c,d**. Stacked barplots show the percentage of alb-like (white), severe (light grey), mild (dark grey), and phenotypically WT (black) embryos 48 hpf after injection. The total number of injected embryos (n) is shown below. The results were obtained from at least two independent experiments. The Fisher test p-value per comparison is shown.

**d.** Schematic representation of *in vivo* experiments in *C. elegans* targeting *dpy-10*. N2 worms are injected with a mixture containing a guide against the *dpy-10* locus. Mutations leading to dominant Rol or Dpy phenotypes are detected in the $F_1$. In some experiments, wild type-like $F_1$ progeny are singled out to screen for recessive Rol or Dpy phenotypes in the $F_2$.

**e.** Comparison of WT Cas9 (WT), SpG, and SpRY editing efficiencies in *C. elegans* using the canonical *dpy-10* co-CRISPR gRNA (Arribere et al., 2014) at an RNP concentration of 1.6 µM. The left panel includes $F_1$ progeny laid between 1 and 24 hr post-injection, the middle panel includes $F_1$ progeny laid after 24 hr post-injection, and the right panel includes all $F_1$ progeny, all from the same set of injected $P_0$s. The editing efficiency is defined as the number of $F_1$ progeny with Rol or Dpy phenotypes divided by the total number of $F_1$ progeny laid by each $P_0$.

Each dot represents the editing efficiency in each individual $P_0$ that produced at least 100 $F_1$. The numbers indicate the total number of viable injected $P_0$s and screened $F_1$s per group. The results are obtained from two independent experiments, with all three conditions carried out in parallel injections. $p<0.0001$ **** (One-way ANOVA followed by Tukey's test for multiple comparisons).

**Supplementary Figure 3. Comparative activities of IDT Cas9, WT SpCas9, SpG, and SpRY against the *dpy-10* NGG target *in vitro.***
Four different RNP combinations comprised of each guide and IDT Cas9, WT SpCas9, SpG, or SpRY were tested *in vitro* at 15, 25, 37, and 50 °C by incubating the RNPs with a PCR product amplified from the *dpy-10* locus for one hour. Negative controls without the guide RNA (-gRNA) were tested at 37 °C. The top row of bands shows uncleaved PCR product at 698 bp, the middle row of bands shows cleaved products at 432 bp, and the bottom row of bands shows cleaved products at 266 bp.

**Supplementary Figure 4. SpG is an efficient nuclease at minimal PAM targets in zebrafish.**
**a-b.** Phenotypic evaluation of injected embryos with different amounts of mRNAs coding SpG (mSpG) and gRNAs (1-5) targeting NGH PAM sites for *slc45a2* (*albino*) (**a**) or *slc24a5* (*golden*) (**b**) genes. Stacked barplots show the percentage of albino/golden-like (white), severe (light grey), mild (dark grey), and phenotypically WT (black) embryos 48 hpf after injection. The total number of injected embryos (n) is shown below. Data in optimal conditions (300 pg mRNA and 240 pg gRNA per embryo) from **Fig. 2b**. The results were obtained from at least two independent experiments.
**c.** Phenotypic evaluation of injected embryos with different amounts of mRNAs coding for SpG (mSpG) and gRNAs (1-5) targeting NGH PAM sites for *tbxta* (*no-tail*) gene. Stacked barplots show the percentage of class III (white), class II (light grey), class I (dark grey), and phenotypically WT (black) embryos 28 hpf after injection. The total number of injected embryos (n) is shown below. Data in optimal conditions (300 pg mRNA and 240 pg gRNA per embryo) from **Fig. 2b**. The results were obtained from at least two independent experiments.

**Supplementary Figure 5. Examples of mutations obtained with SpG complexes targeting *slc45a2, slc24a5* and *tbxta* genes in zebrafish; and endogenous SpG (SpGe) targeting *dpy-10* and WT Cas9, SpG, and SpRY targeting wrmScarlet in *C. elegans*.**
**a**. Five 24 hpf zebrafish embryos per condition were collected for DNA extraction (see Methods). Different targeting (golden 5, albino 1, and no-tail1) regions were PCR amplified and sequenced. The reference sequence (WT) is provided as the first sequence in each group of alignments.; The PAM sequence is underlined; -: deletion.
**b**. Each sequence is derived from individual worms consisting of either $F_2$ dumpies or $F_2$ wrmScarlet knockouts. WT, SpG, and SpRY indicate the nuclease used to edit the genome. SpGe indicates that the animals were edited using the endogenously expressed SpG from the germline. For the *dpy-10* alignments, R indicates that the allele is dominant, and r indicates recessive. The reference sequence (WT) is provided as the first sequence in each group of alignments. The PAM sequence is underlined; -: deletion.

**Supplementary Figure 6. SpRY, but not WT Cas9, is an efficient nuclease at minimal PAM targets in zebrafish**

**a-b.** Phenotypic evaluation of injected embryos with different amounts of mRNAs coding for SpRY (mSpRY) and gRNAs (1-3) targeting NAN PAM sites for *slc45a2* (albino) (**a**) or *slc24a5* (golden) (**b**) genes. Stacked barplots show the percentage of albino/golden-like (white), severe (light grey), mild (dark grey), and phenotypically WT (black) embryos 48 hpf after injection. The total number of injected embryos (n) is shown below. Data in optimal conditions (300 pg mRNA and 240 pg gRNA per embryo) from **Fig. 2f**. The results were obtained from at least two independent experiments.

**c.** Phenotypic evaluation of injected embryos with different amounts of mRNAs coding for SpRY (mSpRY) and gRNAs (1-3) targeting NAN PAM sites for *tbxta* (no-tail) gene. Stacked barplots show the percentage of class III (white), class II (light grey), class I (dark grey), and phenotypically WT (black) embryos 28 hpf after injection. The total number of injected embryos (n) is shown below. Data in optimal conditions (300 pg mRNA and 240 pg gRNA per embryo) from **Fig. 2f**. The results were obtained from at least two independent experiments.

**d.** Phenotypic evaluation of embryos injected with 300 pg of mCas9 and 240 pg of gRNAs that show high activity in targets with NGH and NAN PAM sequence for three genes (*slc45a2*, albino; *slc24a5*, golden; *tbxta*, no-tail, **Fig. 2**) when used with mSpG and mSpRY. Stacked barplots show the percentage of albino/golden-like/class III (white), severe/class II (light grey), mild/class I (dark grey), and phenotypically WT (black) embryos 24 and 48 hpf after injection. The total number of injected embryos (n) is shown below. The results were obtained from at least two independent experiments.

**Supplementary Figure 7. Salt concentration does not affect the editing efficiency of SpG RNP *in vivo***

**a.** Salt concentration does not affect the editing efficiency of an anti-wrmScarlet:SpG RNP (NGH PAM). RNPs consisting of SpG and an anti-wrmScarlet gRNA at the basal 1.3 μM concentration were injected in the CER541 strain to target the wrmScarlet protein with an NGH PAM. Two injection mixes with a final KCl concentration of either 50 mM or 300 mM were compared. The editing efficiency is defined as the number of $F_1$ worms exhibiting loss of fluorescence in the $F_2$ divided by the total number of separated *dpy-10* co-edited $F_1$s. Each dot represents the editing efficiency in each individual $P_0$ that produced at least eight Dpy or Rol $F_1$s. The numbers indicate the total number of viable injected $P_0$s and screened $F_1$s per group. The horizontal black lines represent the median of seven $P_0$s for each salt concentration, and the grey outlines are violin plots. The results were obtained from a single experiment, with both conditions carried out in parallel injections. ns: no significant difference (student's *t*-test)

**b.** Salt concentration does not affect the editing efficiency of a *dpy-10*:SpG RNP (NGG PAM). RNPs consisting of SpG and a *dpy-10* gRNA at the basal 1.3 μM concentration were injected in the N2 strain to target the *dpy-10* gene with an NGG PAM. Two injection mixes with a final KCl concentration of either 50 mM or 300 mM were compared. The editing efficiency is defined as the number of $F_1$ progeny with Rol or Dpy phenotypes divided by the total number of $F_1$ progeny laid by each $P_0$. Each dot represents the editing efficiency in each individual $P_0$ that produced at least 100 $F_1$. The horizontal black lines represent the median of ten to 11 $P_0$s for each salt concentration, and the grey outlines are violin plots. The results were obtained from a single experiment, with both conditions carried out in parallel injections. ns: no significant difference (student's *t*-test).

**Supplementary Figure 8. RNP can increase SpG/SpRY activity in zebrafish.**
**a.** Phenotypic evaluation of injected embryos with 3 or 6 μM (fmol/embryo) of RNP complexes containing Cas9 (WT, grey), SpG (blue), or SpRY (green) proteins and gRNA a and b for *slc45a2* (*albino*) gene. 6 fmol of RNP is equivalent to 1 ng of protein and 240 pg of gRNA per embryo. Stacked barplots show the percentage of albino-like (white), severe (light grey), mild (dark grey), and phenotypically WT (black) embryos 48 hpf after injection. The total number of injected embryos (n) is shown below. The results were obtained from at least two independent experiments.
**b-c.** Comparison of the activity between mSpG and SpG RNP (**b**) or mSpRY and SpRY RNP (**c**) using different gRNAs targeting genomic sites with NGH or NAN PAM sequence respectively in three genes (*slc45a2*, *albino*; *slc24a5*, *golden*; *tbxta*, *no-tail*). Phenotypic evaluation of embryos injected with 300 pg of mSpG/mSpRY and 240 pg of each gRNA or embryos injected with 6 μM (6 fmol/embryo) of RNP complexes containing SpG/SpRY protein and gRNA. mSpG and mSpRY data from **Supplementary Fig. 3 and 6, respectively**. Stacked barplots show the percentage of albino-like/golden-like/class III (white), severe/class II (light grey), mild/class I (dark grey), and phenotypically wild-type (WT; black) embryos 24 and 48 h post fertilization (hpf) after injection. The total number of injected embryos (n) is shown below. The results were obtained from at least two independent experiments.

**a**

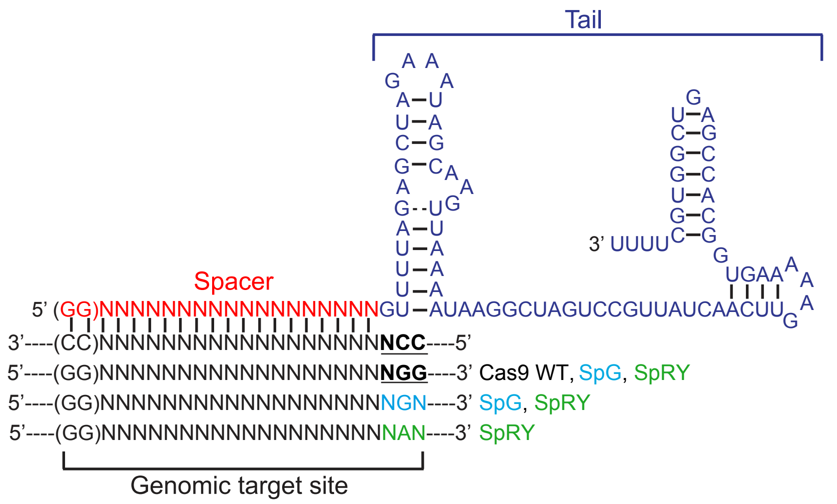
Tail
Spacer
5' (GG)NNNNNNNNNNNNNNNNNNNGU
3'----(CC)NNNNNNNNNNNNNNNNNNNNCC----5'
5'----(GG)NNNNNNNNNNNNNNNNNNNNGG----3' Cas9 WT, SpG, SpRY
5'----(GG)NNNNNNNNNNNNNNNNNNNNGN----3' SpG, SpRY
5'----(GG)NNNNNNNNNNNNNNNNNNNNAN----3' SpRY
Genomic target site
3' UUUU
AUAAGGCUAGUCCGUUAUCAACUU

**b**

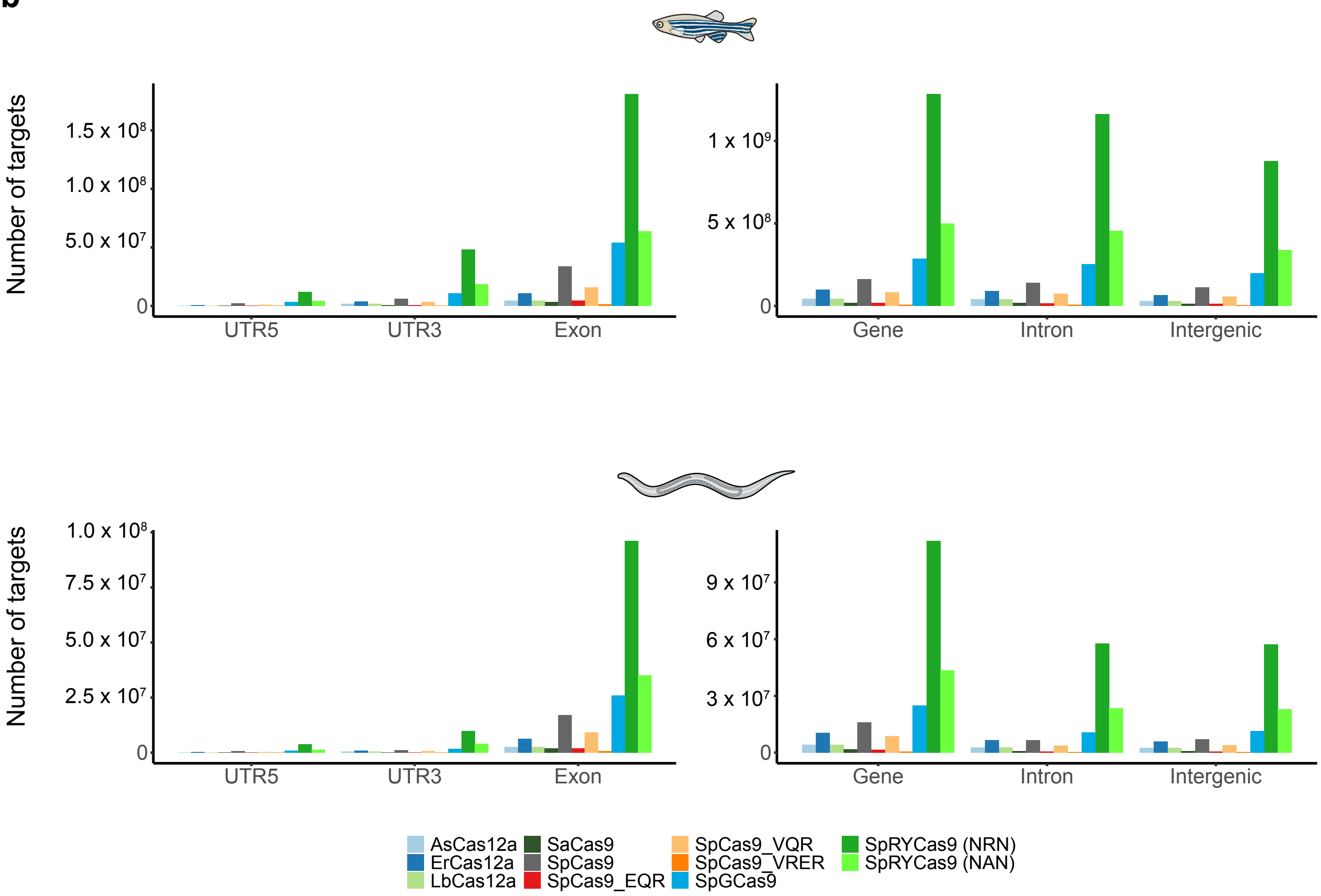
Number of targets
1.5 x 10^8
1.0 x 10^8
5.0 x 10^7
0
UTR5
UTR3
Exon
1 x 10^9
5 x 10^8
Gene
Intron
Intergenic
1.0 x 10^8
7.5 x 10^7
2.5 x 10^7
9 x 10^7
6 x 10^7
3 x 10^7
AsCas12a
ErCas12a
LbCas12a
SaCas9
SpCas9
SpCas9_EQR
SpCas9_VQR
SpCas9_VRER
SpGCas9
SpRYCas9 (NRN)
SpRYCas9 (NAN)

#### Supplementary Figure 2.

Vicencio, Sanchez-Bolaños, et al.

**a**

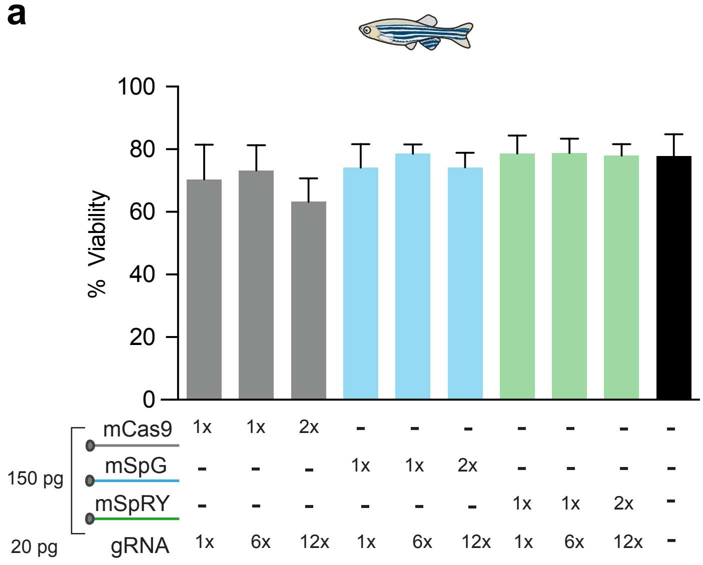


**b**

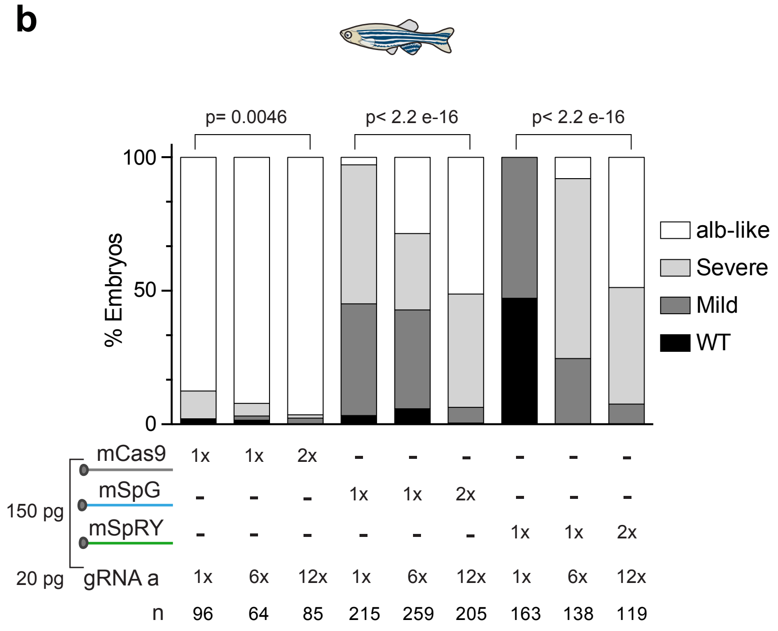


**c**

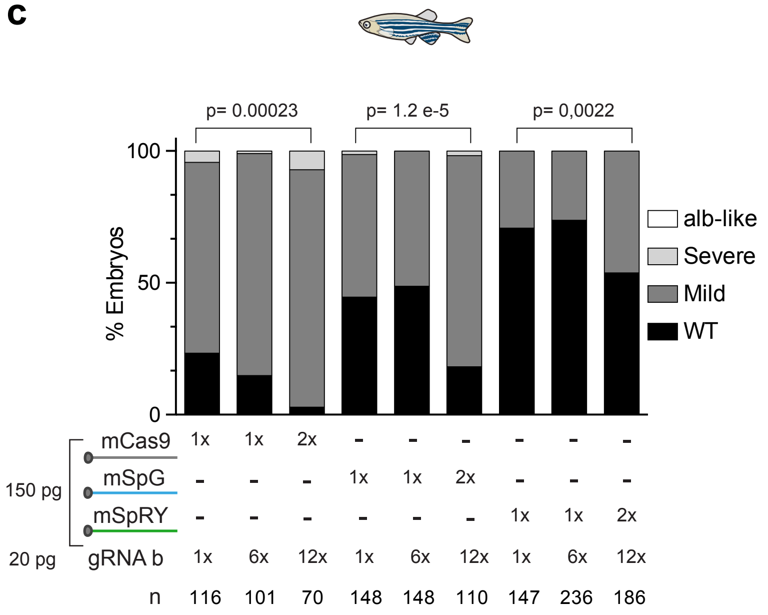


**d**

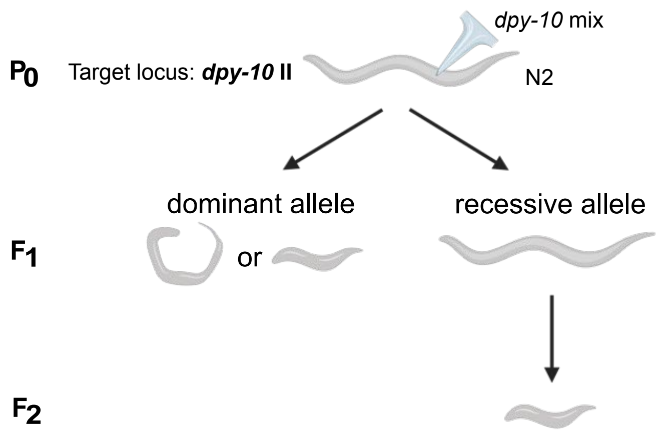


**e**

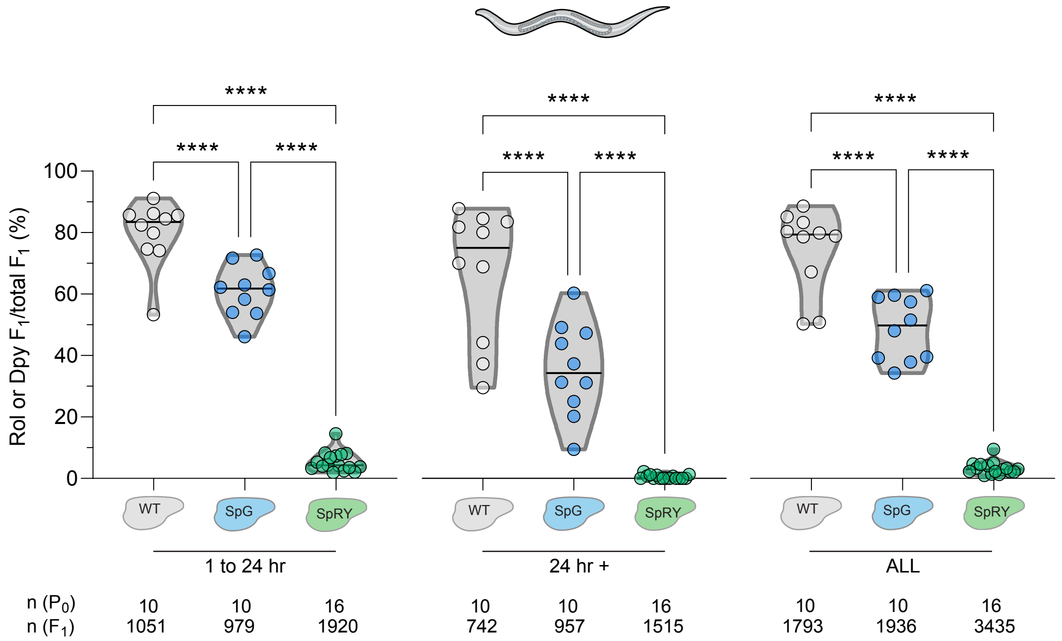

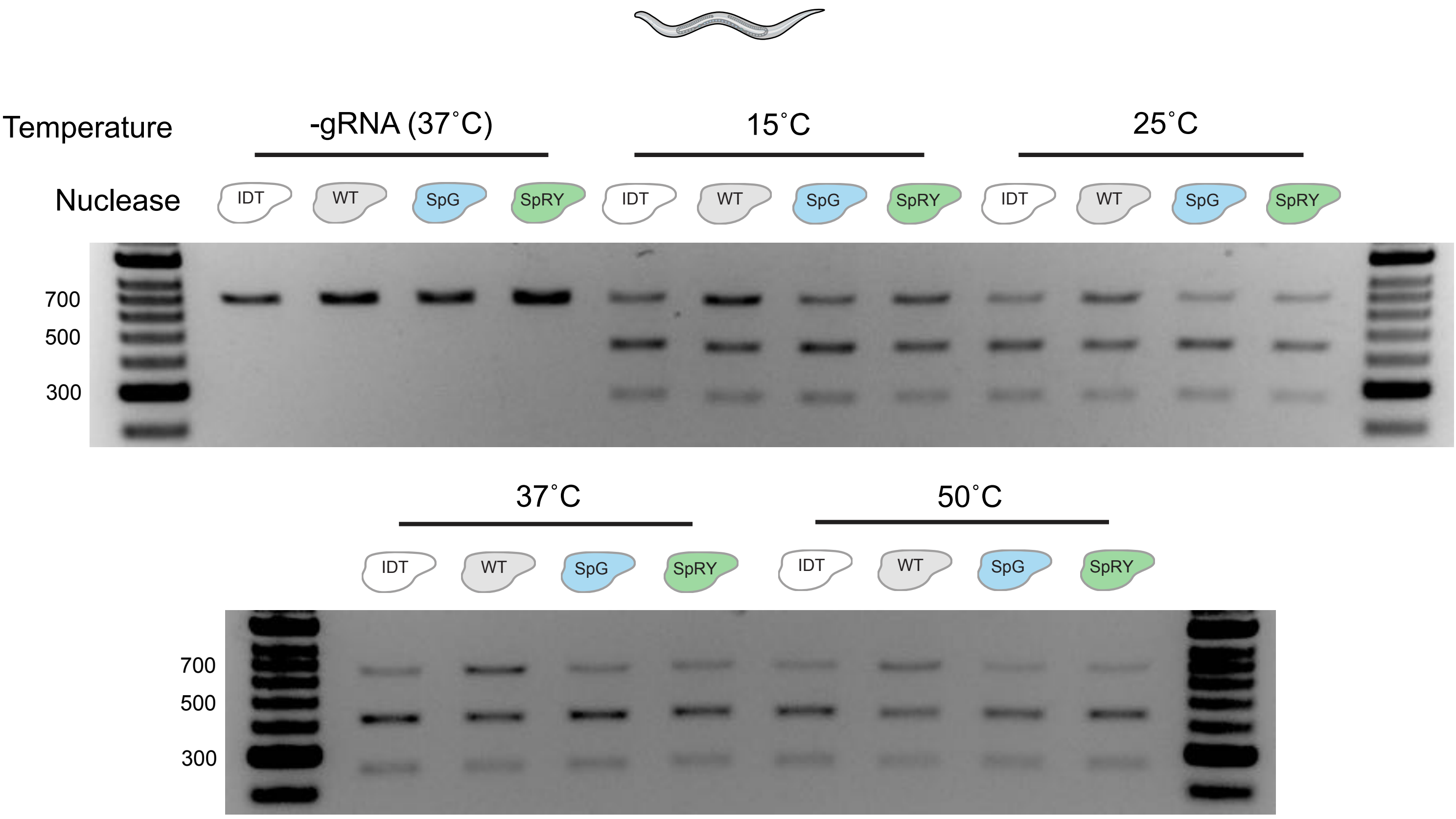
Temperature
-gRNA (37˚C)
15˚C
25˚C
Nuclease
IDT
WT
SpG
SpRY
IDT
WT
SpG
SpRY
IDT
WT
SpG
SpRY
700
500
300
37˚C
50˚C
IDT
WT
SpG
SpRY
IDT
WT
SpG
SpRY
700
500
300

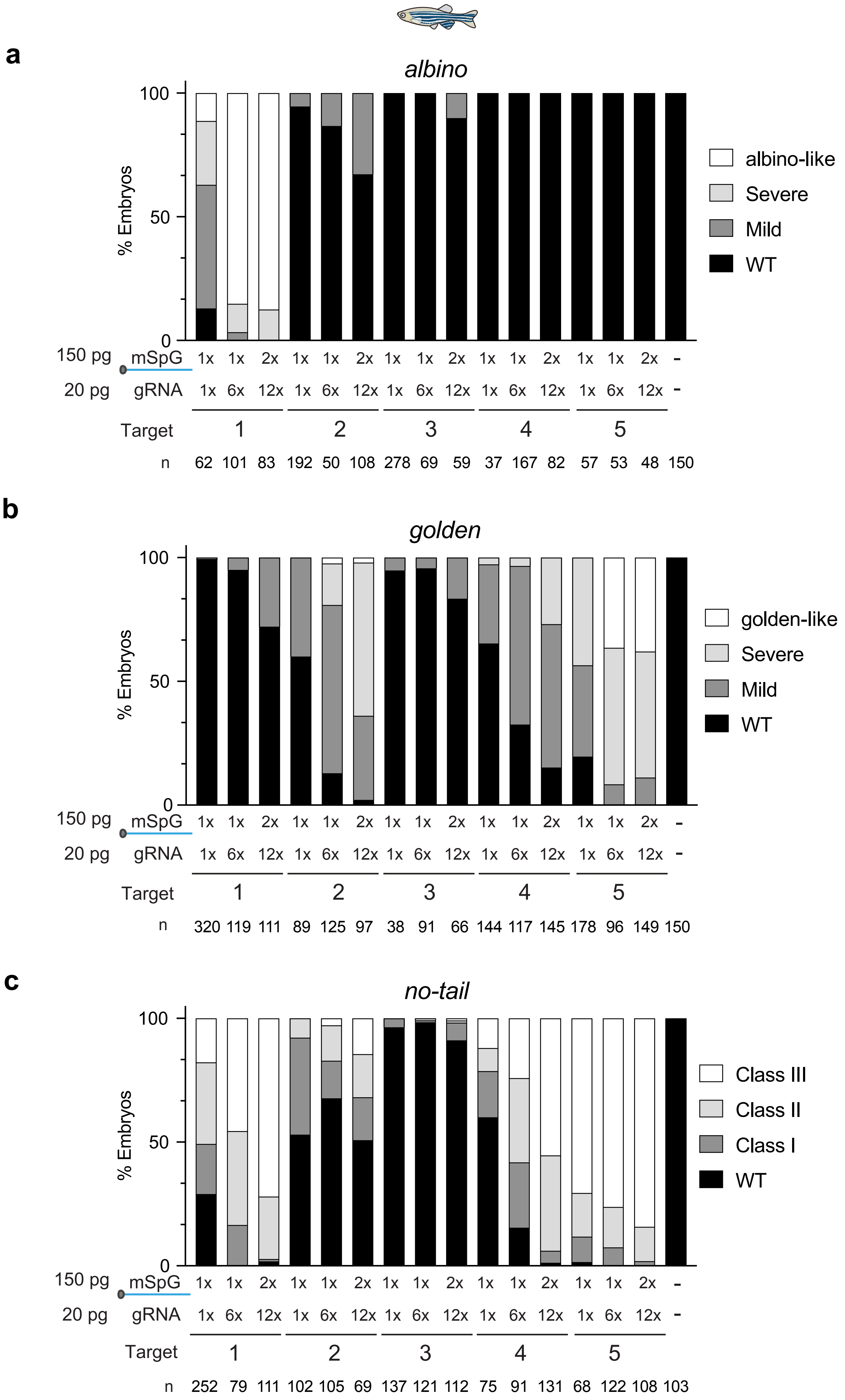
a
albino
100
50
0
% Embryos
albino-like
Severe
Mild
WT
150 pg mSpG 1x 1x 2x 1x 1x 2x 1x 1x 2x 1x 1x 2x 1x 1x 2x -
20 pg gRNA 1x 6x 12x 1x 6x 12x 1x 6x 12x 1x 6x 12x 1x 6x 12x -
Target 1 2 3 4 5
n 62 101 83 192 50 108 278 69 59 37 167 82 57 53 48 150
b
golden
golden-like
Severe
Mild
WT
150 pg mSpG 1x 1x 2x 1x 1x 2x 1x 1x 2x 1x 1x 2x 1x 1x 2x -
20 pg gRNA 1x 6x 12x 1x 6x 12x 1x 6x 12x 1x 6x 12x 1x 6x 12x -
Target 1 2 3 4 5
n 320 119 111 89 125 97 38 91 66 144 117 145 178 96 149 150
c
no-tail
Class III
Class II
Class I
WT
150 pg mSpG 1x 1x 2x 1x 1x 2x 1x 1x 2x 1x 1x 2x 1x 1x 2x -
20 pg gRNA 1x 6x 12x 1x 6x 12x 1x 6x 12x 1x 6x 12x 1x 6x 12x -
Target 1 2 3 4 5
n 252 79 111 102 105 69 137 121 112 75 91 131 68 122 108 103

**Supplementary Figure 5.** **Vicencio, Sanchez-Bolaños, et al.**

**a**

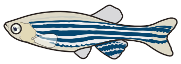

##### Target *golden* 5

```
WT  TGGGTTTCAGGTCTTGGTCTCGCAGGATGTTGCTGGAGCCACGTTTATGGCTGCGGGGAGTTCGGCTCCAGAG
    TGGGTTTCAGGTCTTGGTCTCGC---ATGTTGCTGGAGCCACGTTTATGGCTGCGGGGAGTTCGGCTCCAGAG
    TGGGTTTCAGGTCTTGGTCTCG----ATGTTGCTGGAGCCACGTTTATGGCTGCGGGGAGTTCGGCTCCAGAG
    TGGGTTTCAGGTCTTGGTCTCG-------TTGCTGGAGCCACGTTTATGGCTGCGGGGAGTTCGGCTCCAGAG
    TGGGTTTCAGGTCTTGGTCTCGC-------TGCTGGAGCCACGTTTATGGCTGCGGGGAGTTCGGCTCCAGAG
    TGGGTTTCAGGTCTTGGTCTCGCNAGGATGTTGCTGGAGCCACGTTTATGGCTGCGGGGAGTTCGGCTCCAGA
```

##### Target *albino* 1

```
WT  CTACTGTAGGTCGTCATGGGGCCGAAGGAGACCGTACATACTCTTACTGGGGATTCTGATGTTAGTGGGCATGAC
    CTACTGTAGGTCGTCATGGGGCC----------GAACATACTCTTACTGGGGATTCTGATGTTAGTGGGCATGAC
    CTACTGTAGGTCGTCATGGGGCC----------GTACATACTCTTACTGGGGATTCTGATGTTAGTGGGCATGAC
    CTACTGTAGGTCGTCATGGGGCC---------GATACATACTCTTACTGGGGATTCTGATGTTAGTGGGCATGAC
    CTACTGTAGGTCGTCATGGAGAC---------CGTACATACTCTTACTGGGGATTCTGATGTTAGTGGGCATGAC
    CTACTGTAGGTCGTCATGGGG------------------ACTCTTACTGGGGATTCTGATGTTAGTGGGCATGAC
    CTACTGTAGGTCGTCATGGGGAC---------CGTACATACTCTTACTGGGGATTCTGATGTTAGTGGGCATGAC
    CTACTGTAGGTCGTCATGGGGCC---------GGTACATACTCTTACTGGGGATTCTGATGTTAGTGGGCATGAC
    CTACTGTAGGTCGTCATGGGG----AGGAGACCGTACATACTCTTACTGGGGATTCTGATGTTAGTGGGCATGAC
    CTACTGTAGGTCGTCATGGGGCCGANAGGAGACCGTACATACTCTTACTGGGGATTCTGATGTTAGTGGGCATGA
```

##### Target *no-tail* 1

```
WT  ACGGGTGCTTTCATCCAGTGCGCGCCGAAGTTGGGTGAGTCCGGGTGGATGTAGACGCAGCTCGGGCTTTGGGGT
    ACGGGTGCTTTCATCCAGTGCGCG---AAGTTGGGTGAGTCCGGGTGGATGTAGACGCAGCTCGGGCTTTGGGGT
    ACGGGTGCTTTCATCCAGTGCGCG---CAGTTGGGTGAGTCCGGGTGGATGTAGACGCAGCTCGGGCTTTGGGGT
    ACGGGTGCTTTCATCCAGTGCGCA-----GTTGGGTGAGTCCGGGTGGATGTAGACGCAGCTCGGGCTTTGGGGT
    ACGGGTGCTTTCATCCAGTGCGCG-----GTTGGGTGAGTCCGGGTGGATGTAGACGCAGCTCGGGCTTTGGGGT
    ACGGGTGCTTTCATCCAGTGCGCG-CGAAGTTGGGTGAGTCCGGGTGGATGTAGACGCAGCTCGGGCTTTGGGGT
    ACGGGTGCTTTCATC------------AAGTTGGGTGAGTCCGGGTGGATGTAGACGCAGCTCGGGCTTTGGGGT
    ACGGGTGCTTTCATCCAGTGCGCGCNCGAAGTTGGGTGAGTCCGGGTGGATGTAGACGCAGCTCGGGCTTTGGGG
```

**b**

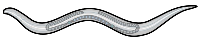

##### Target *dpy-10*

```
WT      CTTCAATACGGCAAGATGAGAATGACTGGAAACCGTACCGCTCGTGGTGCCTATGGTAGCGGAGCTTCACATGGCTTCAG
SpGe R  CTTCAATACGGCAAGATGAGAATGACTGGAAAC---------CGTGGTGCCTATGGTAGCGGAGCTTCACATGGCTTCAG
SpGe R  CTTCAATACGGCAAGATGAGAATGACTGGAAACCG---TACTCGTGGTGCCTATGGTAGCGGAGCTTCACATGGCTTCAG
SpGe R  CTTCAATACGGCAAGATGAGAATGACTGGAAACCG---TACTCGTGGTGCCTATGGTAGCGGAGCTTCACATGGCTTCAG
SpGe r  CTTCAATACGGCAAGATGAGAATGACTGGAAAC-----CGCTCGTGGTGCCTATGGTAGCGGAGCTTCACATGGCTTCAG
SpGe r  CTTCAATACGGCAAGATGAGAATGACTGGAAACT----CGCTCGTGGTGCCTATGGTAGCGGAGCTTCACATGGCTTCAG
SpGe r  CTTCAATACGGCAAGATGAGAATGACTGGAAA---------TCGTGGTGCCTATGGTAGCGGAGCTTCACATGGCTTCAG
```

##### Target *wrmScarlet*-NAC2

```
WT       GGACCACTCCCATTCTCCTGGGACATCCTCTCCCCACAATTCATGTACGGATCCCGTGCCTTCACCAAGCACCCAGCCGA
WT Cas9  GGACCACTCCCATTCTCCTGGGAC--------CCCACAATTCATGTACGGATCCCGTGCCTTCACCAAGCACCCAGCCGA
SpRY     GGACCACTCCCATTCTCCTGGGAC----------CACAATTCATGTACGGATCCCGTGCCTTCACCAAGCACCCAGCCGA
SpRY     GGACCACTCCCATTCTCCTGGGACATCC---TCTCACAATTCATGTACGGATCCCGTGCCTTCACCAAGCACCCAGCCGA
SpRY     GGACCACTCCCATTCTCCTGGGACAT-----CCCCACAATTCATGTACGGATCCCGTGCCTTCACCAAGCACCCAGCCGA
SpRY     GGACCACTCCCATTCTCCTGGGACATCC---TCACA-AATTCATGTACGGATCCCGTGCCTTCACCAAGCACCCAGCCGA
```

##### Target *wrmScarlet*-NAC3

```
WT    ACAAGCAATCCTTCCCAGAGGGATTCAAGTGGGAGCGTGTCATGAACTTCGAGGACGGAGGAGCCGTCACCGTCACCCAAG
SpRY  ACAAGCAATCCTTCCCAGAGGGATTCAAGTGGGAG-----------CTTCGAGGACGGAGGAGCCGTCACCGTCACCCAAG
```

##### Target *wrmScarlet*-NGH1

```
WT    GACCAGTCATGCAAAAGAAGACCATGGGATGGGAGGCCTCCACCGAGCGTCTCTACCCAGAGGACGGAGTCCTCAAGGGAG
SpG   GACCAGTCATGCAAAAGAAGACCATGGGATGGGAG---------GAGCGTCTCTACCCAGAGGACGGAGTCCTCAAGGGAG
SpG   GACCAGTCATGCAAAAGAAGACCATGGGATGGGAG---------GGGCGTCTCTACCCAGAGGACGGAGTCCTCAAGGGAG
SpRY  GACCAGTCATGCAAAAGAAGACCATGGGATGGGAGGCCT---CCGAGCGTCTCTACCCAGAGGACGGAGTCCTCAAGGGAG
SpRY  GACCAGTCATGCAAAAGAAGACCATGGGATGGGAGG------CCGAGCGTCTCTACCCAGAGGACGGAGTCCTCAAGGGAG
```

**Supplementary Figure 6.**

**Vicencio, Sanchez-Bolaños, et al.**

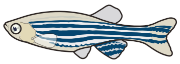

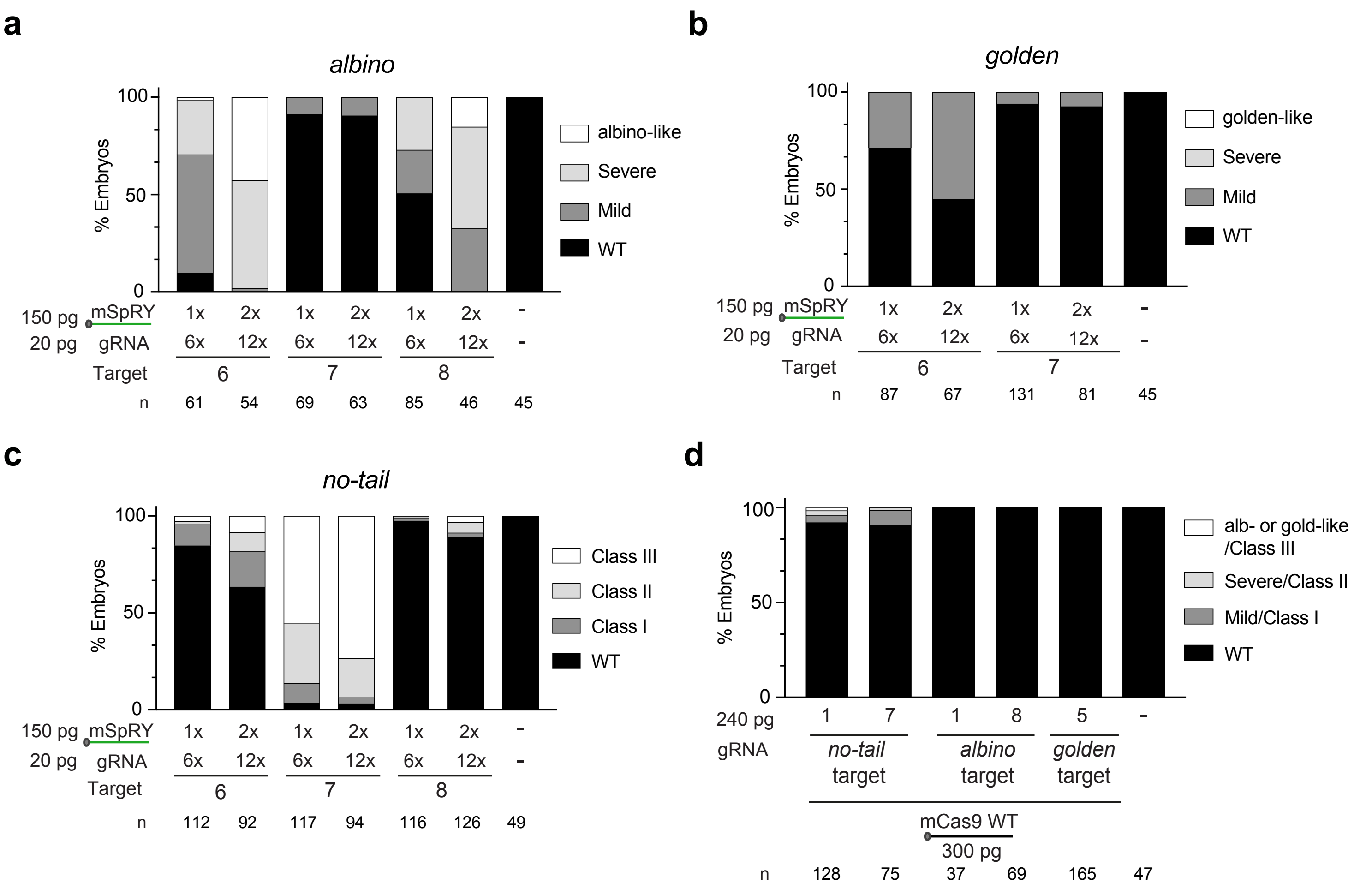

**Supplementary Figure 7.** **Vicencio, Sanchez-Bolaños, et al.**

**a** **b**

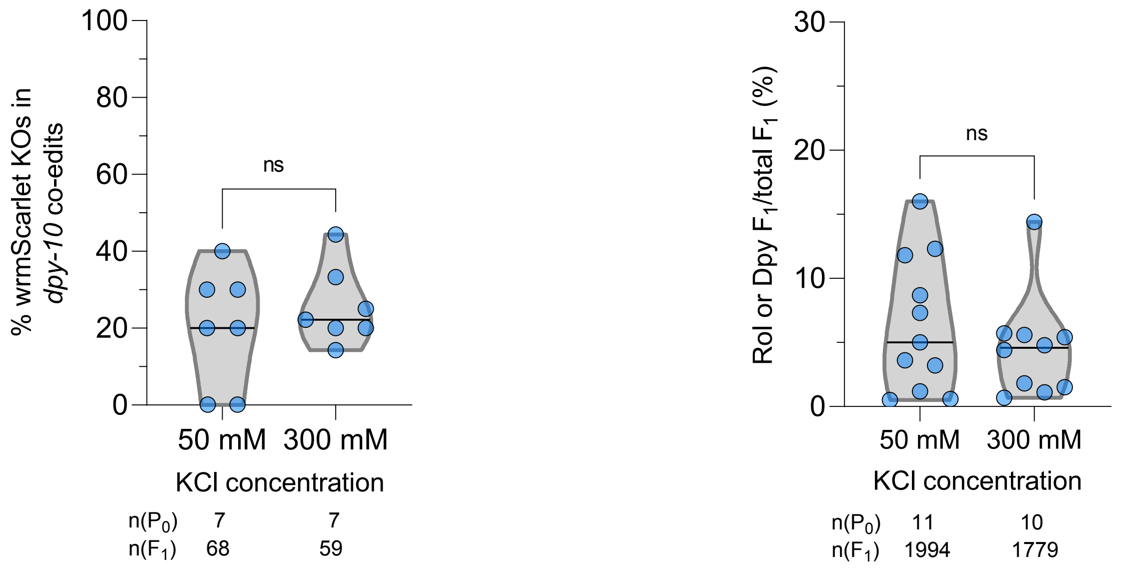

### Supplementary Figure 8.

Vicencio, Sanchez-Bolaños, et al.

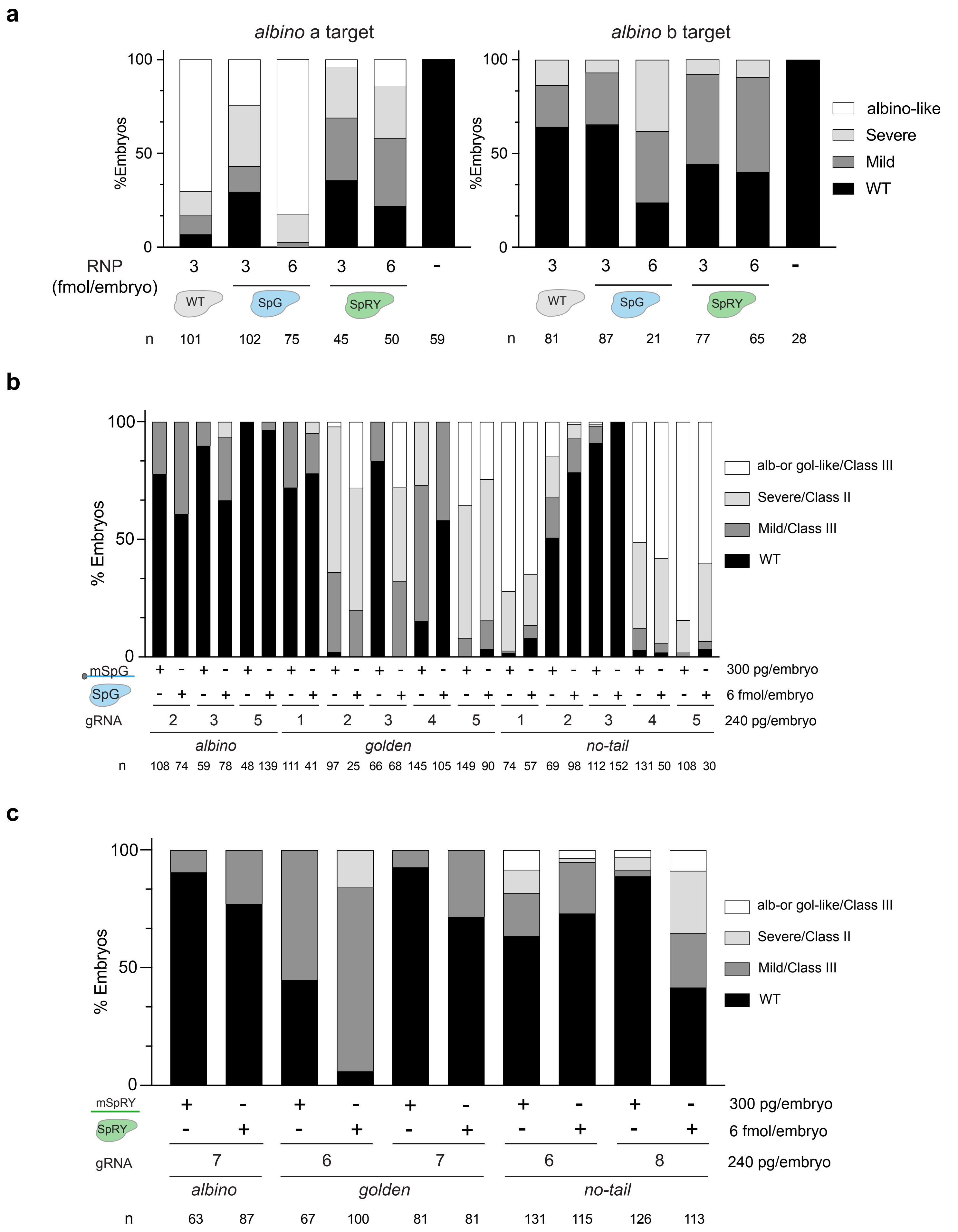
